# Integrative Spatial Analysis of H&E and IHC Images Identifies Prognostic Immune Subtypes Correlated with Progression-Free Survival in Human Papillomavirus (HPV)-Related Oropharyngeal Squamous Cell Carcinoma

**DOI:** 10.1101/2023.09.13.557587

**Authors:** Sumanth Reddy Nakkireddy, Inyeop Jang, Minji Kim, Linda X. Yin, Michael Rivera, Joaquin J. Garcia, Kathleen R. Bartemes, David M. Routman, Eric. J. Moore, Chadi N. Abdel-Halim, Daniel J. Ma, Kathryn M. Van Abel, Tae Hyun Hwang

**Author notes:** **CORRESPONDING AUTHOR:** Tae Hyun Hwang, PhD, Florida Department of Health Cancer Chair, Dept. of AI and Informatics, Cancer Biology and Immunology, Mayo Clinic, 4500 San Pablo Rd S, Jacksonville, FL, 32224. Co-First authors (Sumanth Reddy Nakkireddy, Inyeop Jang, Minji Kim). Co-Senior authors (Daniel J. Ma, Kathryn M. Van Abel, Tae Hyun Hwang). **DISCLOSURES:** None. **FUNDING:** Internal only. **AUTHOR CONTRIBUTIONS:** Sumanth Reddy N: conception, data interpretation, manuscript writing, and revision; Inyeop Jang: data interpretation, manuscript writing, and revision; Minji Kim: data interpretation, manuscript revision; Linda X. Yin: data acquisition; Michael Rivera: study design, data acquisition (pathology review); Joaquin Garcia: study design, data acquisition (pathology review); Kathleen Bartemes: conception, study design, data acquisition, manuscript revision; David Routman: conception, study design, data interpretation, manuscript revision; Daniel Ma: Data interpretation, manuscript revision; Eric Moore: data interpretation, manuscript revision; Chadi A. Halim: data acquisition; Kathryn Van Abel: conception, study design, data acquisition, data interpretation, manuscript writing, and revision; Tae Hyun Hwang: conception, study design, data acquisition, data interpretation, manuscript writing and revision. **Translational Relevance** Although the majority of HPV (+) oropharyngeal squamous cell carcinoma (OPSCC) patients exhibit a favorable prognosis, approximately 20% face recurrent or metastatic disease, posing management challenges. Therefore, identification of robust prognostic markers for risk stratification is essential. Our study focused on the comprehensive characterization of intratumor heterogeneity within the tumor immune microenvironment (TME) in both primary tumors and metastatic nodes. Utilizing computational approaches, we integrated hematoxylin and eosin (H&E) and adjacent immunohistochemistry (IHC)- stained slides to investigate the cellular composition and functional characteristics across different regions within the TME. Based on these detailed immune characteristics, we classified patients into specific immune subtypes. Our TME analysis indicated that patients with high tumor-infiltrating lymphocytes (TIL), increased CD8+ levels, and reduced CD163+ cell counts within their primary tumors were likely to have a more favorable progression-free survival outcome.

## Abstract

**Purpose:** Deep learning techniques excel at identifying tumor-infiltrating lymphocytes (TILs) and immune phenotypes in hematoxylin and eosin (H&E)-stained slides. However, their ability to elucidate detailed functional characteristics of diverse cellular phenotypes within tumor immune microenvironment (TME) is limited. We aimed to enhance our understanding of cellular composition and functional characteristics across TME regions and improve patient stratification by integrating H&E with adjacent immunohistochemistry (IHC) images.

**Methods:** A retrospective study was conducted on patients with HPV(+) oropharyngeal squamous cell carcinoma (OPSCC). Using paired H&E and IHC slides for 11 proteins, a DL pipeline was used to quantify tumor, stroma, and TILs in the TME. Patients were classified into immune inflamed (IN), immune excluded (IE), or immune desert (ID) phenotypes. By registering the IHC and H&E slides, we integrated IHC data to capture protein expression in the corresponding tumor regions. We further stratified patients into specific immune subtypes, such as IN, with increased or reduced CD8+ cells, based on the abundance of these proteins. This characterization provided functional insight into the H&E-based subtypes.

**Results:** Analysis of 88 primary tumors and 70 involved lymph node tissue images revealed an improved prognosis in patients classified as IN in primary tumors with high CD8 and low CD163 expression (p = 0.007). Multivariate Cox regression analysis confirmed a significantly better prognosis for these subtypes.

**Conclusions:** Integrating H&E and IHC data enhances the functional characterization of immune phenotypes of the TME with biological interpretability, and improves patient stratification in HPV(+) OPSCC.

## Introduction

As cells undergo malignant transformation, the host immune system plays a vital role in orchestrating multifaceted responses [1]. The adaptive immune system exhibits remarkable capabilities in targeting and eliminating cancer cells by creating a proinflammatory tumor microenvironment [2]. This immune response is mediated by various immune cells, including B cells, helper T cells, and cytotoxic T cells, collectively referred to as tumor-infiltrating lymphocytes (TILs) [2,3]. Through their collaborative efforts, these immune cells collaborate to suppress tumor growth and inhibit disease progression.

Within the context of HPV(+) OPSCC, a virogenic disease that develops immune tolerance to persistent HPV infection, the interplay between host adaptive immunity and tumor progression becomes increasingly complex [3]. Extensive research has demonstrated that a higher density of both tumoral and stromal TILs in HPV(+) OPSCC is associated with a better prognosis than their HPV(-) counterparts [4]. This finding indicates a pivotal role of TILs in shaping the clinical outcomes of patients with HPV(+) OPSCC.

While the majority of HPV(+) OPSCC patients exhibit favorable clinical outcomes, up to 20% of patients may experience recurrent or metastatic disease that can be challenging to manage [5, 6]. Consequently, there is an urgent need to identify robust prognostic factors that can effectively stratify high-risk HPV(+) OPSCC patients from low-risk ones, particularly as the field enters an era of treatment de-escalation [7]. Such stratification is essential to optimize treatment strategies and tailor interventions based on individual risk profiles.

In the context of head and neck cancers, research into the prognostic and therapeutic roles of TILs is still in its infancy. Spector et al. demonstrated that a lower CD4 TILs count in pretreatment biopsies of head and neck cancers was associated with decreased overall survival [8]. This finding suggests that the presence and composition of TILs may have implications for disease outcomes in this setting. Wansom et al. proposed that the degree of TIL infiltration does not appear to be related to HPV status and that the association between TIL density and survival is independent of HPV status [9]. Conversely, Nasman et al. found that HPV(+) OPSCCs exhibited a higher density of CD8 and Foxp3 TILs than HPV(-) OPSCCs, which correlated with a better clinical outcome in both HPV(+) and HPV(-) tumors [10].

Previous studies examining the prognostic role of TILs in HPV(+) OPSCC have yielded mixed results, contributing to the ongoing debate surrounding their significance in predicting clinical outcomes [11, 12]. While some studies have reported a significant association between TILs and survival in HPV(+) OPSCC, others suggest that TIL density may be prognostic only in lower-stage disease [12]. However, most of these TIL-based analyses rely on manual quantification performed by pathologists, which introduces inherent inter-observer variability even among experts [13] and is prohibitively time-consuming for clinical use.

Advancements in digital pathology and deep learning (DL) algorithms have paved the way for automated TIL quantification and analysis from pathology slides [12]. DL-based whole-slide hematoxylin and eosin (H&E) image analyses offer a cost-effective and readily applicable approach to quantifying TILs within tumors [12, 14]. H&E-based evaluation of TILs provides information on the presence of specific cell types within the TME but is limited in the evaluation of the functional roles and characteristics of these cells.

In contrast, immunohistochemical (IHC) staining techniques enable the identification and characterization of specific protein markers, illuminating the functional roles of cells within the TME. Integrating the information obtained from both H&E and IHC data can provide more accurate identification of cells, including TILs, in the TME and offer a comprehensive understanding of the functional roles played by various cell populations within the TME.

Motivated by these considerations, our study aimed to develop a comprehensive computational pipeline that integrates paired H&E- and IHC-stained images from primary tumors and matched pathologically involved (pN+) lymph nodes. By leveraging this pipeline, we sought to investigate the functional characteristics and spatial interplay of TILs within different regions of the tissue, such as the tumor and stroma, using H&E images, as well as the cell-level protein expression patterns obtained from registered IHC slides with H&E, to further understand the functional roles of cells within the TME using DL-based analyses.

## Materials and Methods

### Patient cohort

The methodology for the identification and selection of this matched case-control cohort has been previously published [13]. After obtaining institutional review board approval (IRB 20-012036), we queried a departmental REDCap oropharyngeal database to identify patients with HPV(+) OPSCC of the tonsil or base of the tongue who underwent intent-to-cure surgery +/- adjuvant therapy between 05/2007 and 12/2016. Cases developed locoregional or distant recurrences. Controls were matched based on age, sex, pathologic T, N, overall stage, year of surgery, type of adjuvant treatment received, and the Adult Comorbidity Evaluation-27 (ACE-27) score (Table 1). Our patient cohort consisted of patients with known primary tumors (N=88) and pN+ lymph node samples (N=70).

**Table 1.**
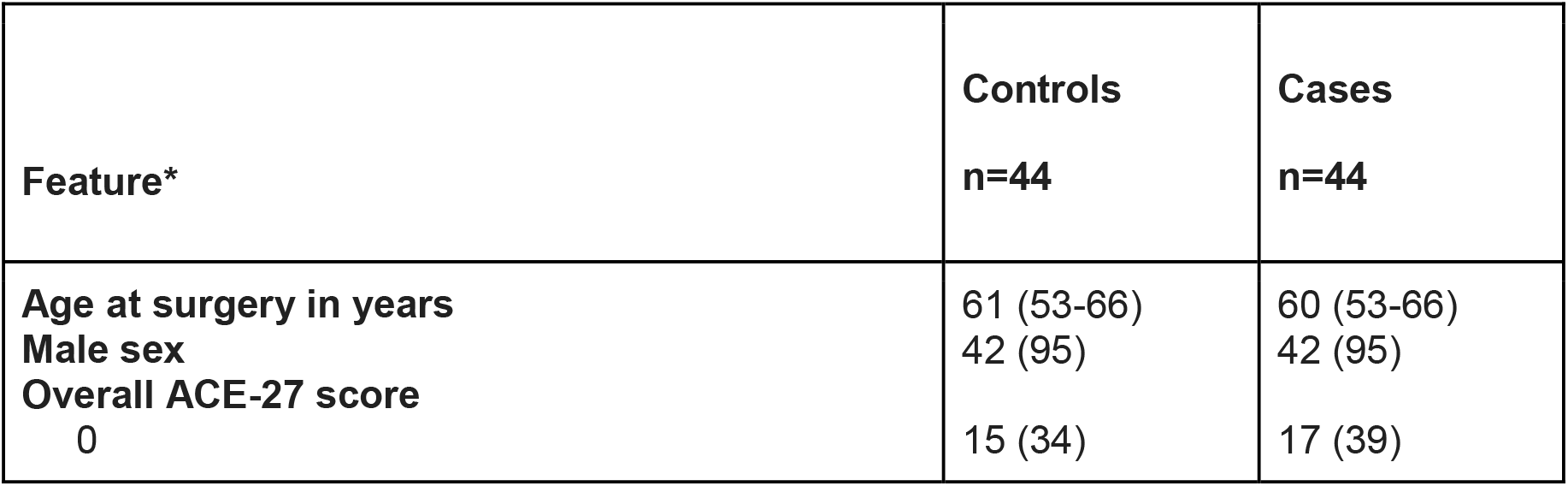
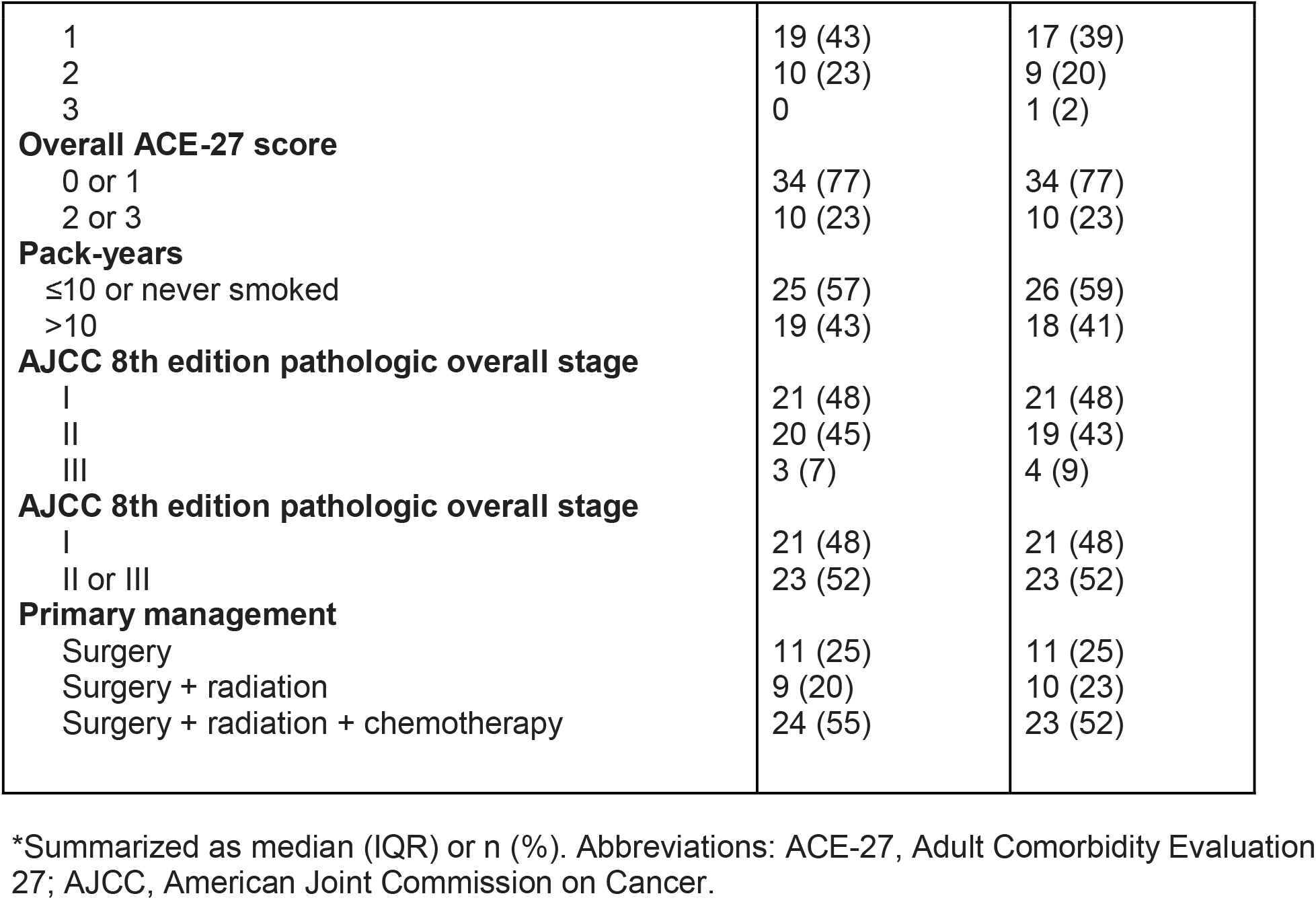
Summary of features used for matching for cases who experienced disease progression and matched controls who did not.

Eligible patients underwent margin-negative transoral resection of the primary tumor with concurrent neck dissection following previously described methods [15]. Exclusion criteria included a history of head and neck cancer, synchronous primary solid tumors in the oropharynx or elsewhere in the body, unknown primary disease (T0), metastatic disease at presentation, and participation in institutional adjuvant radiotherapy de-escalation clinical trials [16].

Within the selected patient cohort meeting the inclusion and exclusion criteria, 44 patients who experienced local, regional, or distant recurrence during the follow-up period and 44 matched controls were identified for analysis. The sample set consisted of 88 diagnostic whole-slide images (WSIs) of primary tumor tissue and 70 images of matched pN+ lymph nodes. H&E and IHC staining were performed on serial tissue sections. Additional clinical data, including the American Joint Commission on Cancer (AJCC) 8th edition pathologic T, N, and overall stage (exact match), sex (exact match), year of surgery (+/- 2 years), type of adjuvant treatment received (none, radiotherapy, or chemoradiotherapy), and Adult Comorbidity Evaluation-27 (ACE-27) score. IHC data for each patient included staining for CD3, CD4, CD8, FoxP3, CD163, ER-alpha, KRT AE1AE3, PD-L1, CD20, CD45-pan, and ER-beta on serial sections of the tissue.

### Pathology review

Formalin-fixed paraffin-embedded (FFPE) tissue specimens were obtained through pathology requisition for all patients (N=88). Two head and neck trained pathologists (MR, JJG) screened all available slides to select one representative H&E slide from the primary tumor and, if available, the metastatic pN+ lymph node for each patient. An Independent blinded review by the two pathologists was conducted to assess TILs density. Density scoring was performed at 100-400x magnification. In cases where the pathologists disagreed on a specific score during independent grading, the slides were reviewed together, and consensus was reached on a final agreed-upon score. Tumoral TILs were defined as lymphocytes directly in contact with tumor cells without intervening stroma. The density was scored as a percentage of the total cell population, categorized as <10%, 10-39%, or ≥40%.

### Immune phenotype classification

We developed a DL-based pipeline (Figure 1A) for spatial analysis of heterogeneous TIL distribution in H&E-stained tissues of different sizes. WSIs were roughly divided into 0.5 mm^2^ grids, and the immune phenotype (IP) of each grid was classified based on the proportion of each component (Figure 2A). Each 0.5 mm^2^ grid was further subdivided into tiles, with each tile containing 112 112 pixels. Each pixel was 0.26μm. Consequently, within each 0.5 mm^2^ grid, there were approximately 17 17 tiles (Figure 2A). The tiles were classified into three categories: TIL, tumors, and stroma using an in-house Resnet18-based pre-trained deep learning model [14].

**Figure 1.**
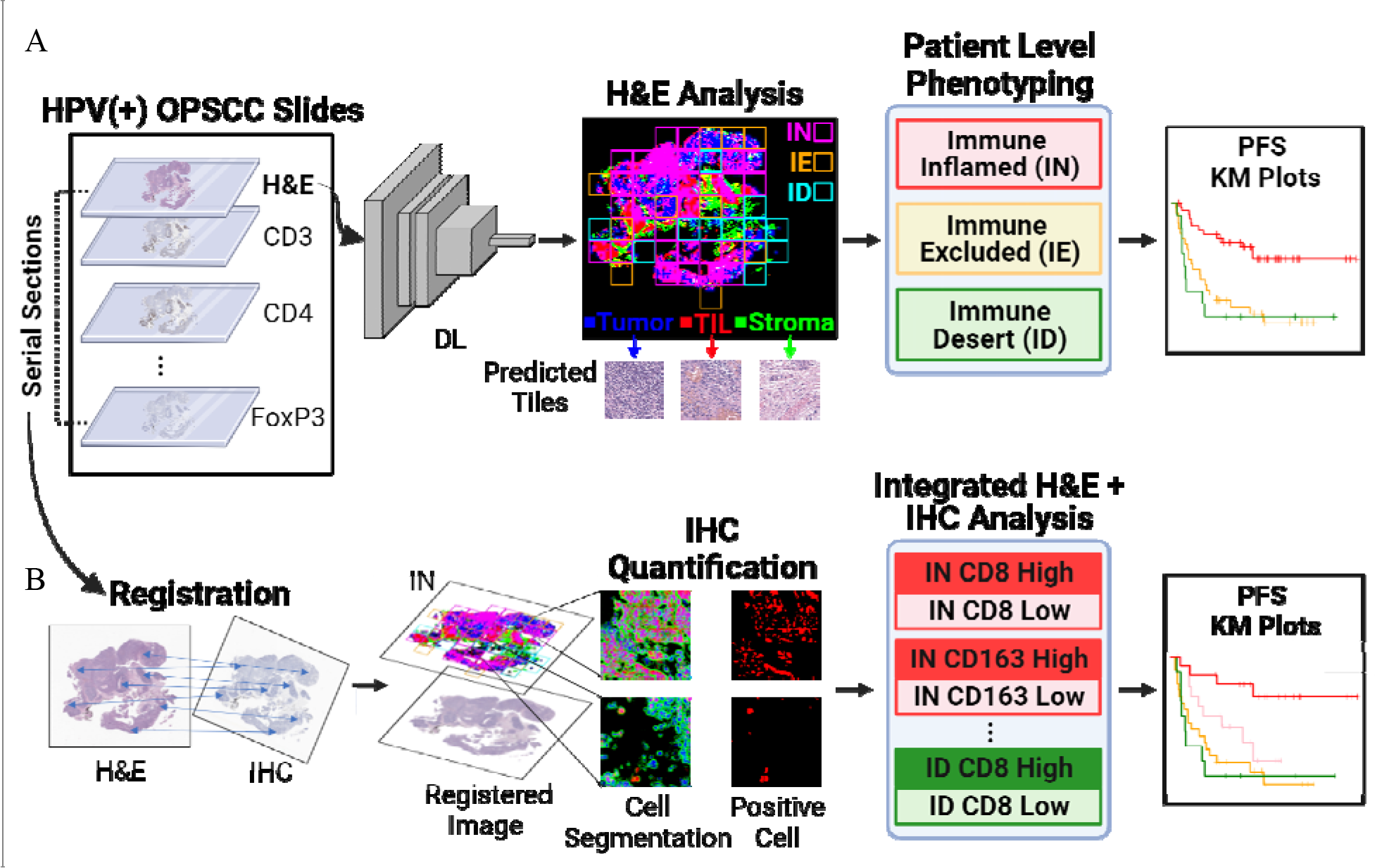
Development of AI-based phenotyping based on integration of both H&E and IHC data. (A) DL-based prediction of the patches in H&E: TIL (red), tumor (blue), stroma (green) and immune phenotyping of 0.5mm^2^ grids based on TIL density: IN (pink), IE (orange), ID (cyan), (middle panel). Patient-level phenotyping is based on the proportion of respective grids followed by survival analysis (right panels). (B) Registration of H&E with IHC panels. Further stratification of IN group patients based on marker expression followed by survival analysis (right panel). TIL, tumor-infiltrating lymphocytes; IN, immune inflamed; IE, immune excluded; ID, immune desert; IHC, immunohistochemistry; H&E, hematoxylin-eosin.

**Figure 2.**
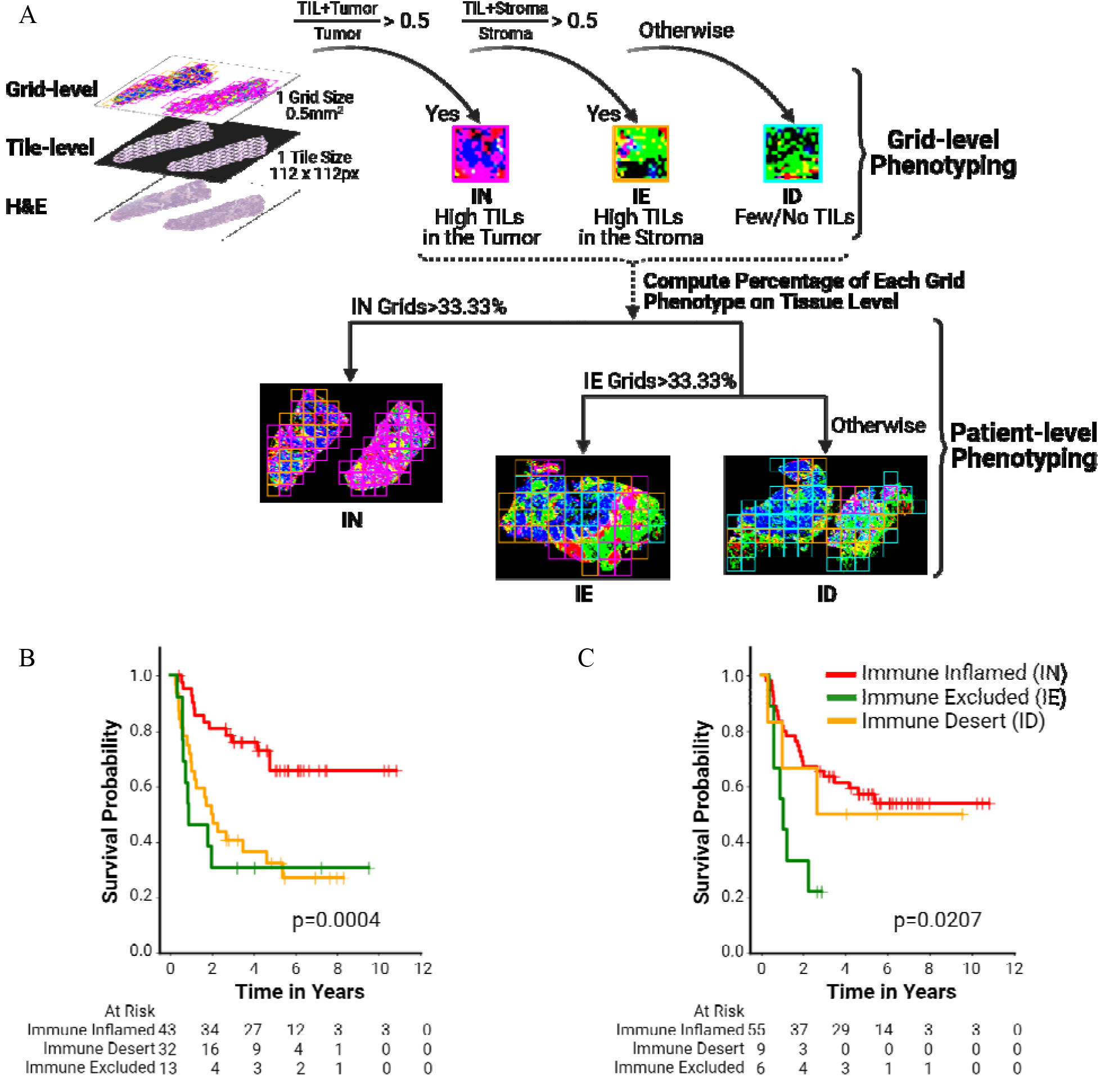
(A) Examples of patient-level phenotyping based on the corresponding 0.5mm^2^ grid phenotyping (top) and thresholding scheme (bottom). (B and C) Kaplan-Meier analysis of progression-free survival based on H&E analysis for primary tumor (B) and pN+ lymph node (C) tissue. IN, immune inflamed; ID, immune desert; IE, immune excluded.

The IP of each 0.5 mm^2^ grid was defined as follows: immune-inflamed (IN) when TIL density in the cancer epithelium (CE) area was above the threshold (>50% tiles); immune-excluded (IE) when TIL density in the CE area was below the threshold and TIL density in the cancer stroma (CS) area was above the threshold (>50% tiles); and immune-desert when TIL density in both the CE area and in the CS area was below the threshold (Figure 2A), similar to the strategy described by Park et al. [12]. The Kaplan-Meier (KM) analysis was carried out with various thresholding strategies; however, the 50% threshold provided a representative separation between different IPs (Supplementary Figure S1). The IN, IE, and ID scores of a WSI were defined by the number of grids annotated to a certain IP divided by the total number of grids analyzed in the WSI. The representative IP of a WSI was defined as the IN phenotype if the IN score was above 33.3%, or IE phenotype if the IE score was above 33.3%, the IN score was below 33.3%, and the ID phenotype otherwise [12].

Quantification of various proteins of interest from the available panel was performed through independent analysis of the IHC slides. To integrate the H&E and IHC data obtained from the serial sections, an image-based registration process was implemented for each patient. Subsequently, the immune subgroups derived from H&E analysis were further stratified based on protein expression levels. Initially, the coordinates of the registered H&E image on a 0.5 mm^2^ grid were identified on the corresponding IHC image. The DeepLIIF algorithm [17] was then employed to identify the number of cells that showed positive staining for the marker of interest within each grid. Using median thresholding, the patients’ H&E-based subgroups were further divided into high and low marker groups based on the quantified number of positive cells for the specific marker of interest in the corresponding grids.

### H&E and IHC image registration

Joint analysis of multiple protein markers and cellular morphology is essential for monitoring multimodal molecular and potentially functional information of cells. To perform the morphological analysis of TME from H&E and the quantitative analysis of proteins from IHC, we used a novel image registration technique. It consists of two steps: rigid and non-rigid registration. In the rigid registration, we adopted a coarse-to-fine transformation to cover both small and large transforms. That is, each pair of H&E and IHC images was resized at 1.0, 0.5, and 0.25 magnification, respectively, to generate an image pyramid of H&E-IHC pairs. At each level, the corresponding pixels between H&E and IHC staining were matched. Next, an initial rigid transformation was performed at the smallest level, the initial transformation was further adjusted by pixel pairs at the intermediate level, and the final adjustment was performed once again in the same manner at the last level.

To account for irregular deformations caused by tissue movement during staining, we applied B-spline-based nonrigid registration [18]. The algorithm uses the previously matched pixel points to control the overall non-rigid deformation and smoothly deforms the pixels between the control points according to the blending coefficients of the spline curve. Finally, we created co-registered IHC images of different markers toward the common H&E image on the serial section (Figure 1A).

### IHC quantification

For the quantitative analysis of proteins, we employed a generative deep neural network, DeepLIIF, that segments cells into a non-nuclear marker (i.e., Ki67), nuclear (DAPI), and nuclear envelope (Lab2) from an IHC image [19]. In particular, we exploited its ability to accurately classify positive or negative cells at the pixel level in the IHC image.

The segmented image from IHC has low contrast and noise pixels, and an automated series of post-processing steps was adopted to account for this. First, a histogram equalizer is employed to improve the contrast by adjusting the pixel intensities of the segmentation image predicted by DeepLIIF. Next, morphological operations (i.e., closing and opening) are applied to the contrast-improved segmentation image to remove tiny groups of pixels or isolated pixels and extract the main components representing each cell. Next, color segmentation is applied to delimit the region of each cell and render them countable. DeepLIIF colored cell boundaries in green, positive cells in red, and negative cells in blue with different intensities (Figure 1B). As each cell had different intensities and unclear boundaries, it was difficult to define the exact region of each cell using a specific color. To overcome this drawback, we converted the color space of the image from RGB to hue, saturation, and value (HSV). In the HSV space, the cell boundaries are much more localized and visually separable. The saturation and value of the color varied, but were mostly located within a small range along the hue axis. Subsequently, the pixels for the boundary were used to detect contours, and only the pixels surrounding the contours were counted as one cell.

### Statistical analysis

The KM method was used to estimate progression-free survival (PFS) in association with the identified immune phenotypes. Hazard ratios (HR) and 95% confidence intervals (CIs) were computed using the Cox proportional hazards model, and a log-rank test was used to assess the level of significant differences between groups in PFS. Two-tailed tests with p-values <0.05 were considered statistically significant. Specimens with inadequate registration of H&E and IHC were excluded from the integrated and Cox regression analyses.

## Data availability

The data generated in this study are available in the article and its supplementary files. The raw data behind the figures and supplementary figures are available upon request from the corresponding authors.

## Code availability

The scripts used for this study are available here: [https://github.com/hwanglab/HE_IHC_HN_analysis]

## Results

### Deep learning-based tissue segmentation for immune phenotype classification

We developed a computational pipeline to integrate H&E staining (N=88) and IHCs to identify immune-based patient subgroups with distinct clinical outcomes (Figure 1). Briefly, we first applied a DL-based algorithm to H&E images to identify the tumor, stroma, and TIL regions within the tumor. The accuracy of the segmentation process for identifying tumors, TILs, and stroma regions based on H&E images has been previously published [14].

Based on the spatial distribution of TILs and stroma within the tumor and other tissue regions, we classified the samples into three distinct immune phenotypes (IPs): IN (high infiltration of TILs in the tumor region), IE (TILs are mostly localized in stroma), and ID (lack of immune cells) [12]. We performed KM analysis to evaluate any statistically significant differences in PFS among the identified subgroups.

For primary tumors, the proportions of patients classified as IN, IE, and ID were 48.9%, 14.8%, and 36.4%, respectively. Interestingly, the IN subgroup showed the most favorable prognosis in both tissue types (log-rank test, p = 0.0004 and 0.0207 in the primary tumor and pN+ lymph nodes, respectively) (Figure 2B and 2C). For example, in the primary tissue, the IN subgroup showed significantly better PFS outcomes than the ID and IE subgroups (Figure 2B). However, in pN+ lymph node tissues, while the IN subgroup showed a statistically significant improvement in PFS compared to the IE subgroup (log-rank test p = 0.004), there was no statistically significant difference between the IN and ID subgroups (log-rank test p-value, 0.67). This discrepancy in PFS between primary tumor and pN+ lymph node tissues suggests that the prognostic utility of TIL density may be tissue-dependent. Specifically, while TIL density may provide prognostic information on PFS in primary tumor tissues, its prognostic significance in pN+ lymph node tissues may be modulated by other factors.

Next, we performed an integrative analysis of the immune phenotypes of the primary tumor and pN+ lymph node data. We found that 30 patients consistently exhibited membership in the IN subgroup associated with a better prognosis (Supplementary Figure S2). The PFS at 1 year was 95.2% for patients with a primary tumor classification of IN and 83.6% with a pN+ lymph node classification of IN. In contrast, the PFS at 1 year was 68.8% and 46.2% for patients with ID and IE phenotypes of the primary tumor and 55.6% and 66.7% with ID and IE of the pN+ lymph node, respectively. By integrating the primary tumor and pN+ lymph node immune phenotypes, we were able to improve the discrimination among the three distinct phenotypes (Supplementary Figure S2).

In addition, we conducted Chi-squared tests to investigate the association between immune subtypes and patient outcomes, categorizing patients with locoregional or distant recurrence as ‘progressors’ and those without such recurrences as ‘non-progressors’. Analysis of primary tumor tissue images revealed a significant association between immune subtypes and both ‘progressors’ and ‘non-progressors’ (p = 0.001) (Supplementary Figure S3). In contrast, the immune subtypes in pN+ lymph nodes did not show a statistically significant association with disease progression or ‘non-progressor’ status (chi-squared test p-value, 0.614) (Supplementary Figure S3). This discrepancy in the association with immune subtypes and ‘progressors’ and ‘non-progressors’ status across primary tumor and pN+ lymph node tissues further indicate the tissue-dependent prognostic factors influencing PFS in HPV(+) OPSCC.

### Comparison of manual annotation and DL approach for assessing TIL density

We compared manual annotation and DL for TILs density assessment. In our previous study, manual annotation categorized the TIL density into three groups: <10%, 10-39%, and ≥40% [13]. We applied the same classification criteria for the DL approach to calculate the ratio of TILs to tumor tiles to measure TIL density. KM analysis demonstrated a significant association between higher tumoral TIL density as assessed by DL and improved PFS (log rank test p-value <0.001, and 0.013, in primary tumors and pN+ lymph nodes, respectively) (Supplementary Figure S4).

Notably, a substantial disparity was observed between patient subgroups based on DL-predicted TIL density and those based on manual annotation. While manual annotation identified only one patient in the pN+ lymph nodes subgroup and three patients in the primary tumor subgroup with a ≥40% TIL density, DL prediction identified 47 patients in each subgroup. Furthermore, closer examination revealed that 30 patients consistently fell within the ≥40% TIL density subgroup when assessed using DL, combining pN+ lymph node and primary tumor scores. In addition, the DL approach identified a substantially higher number of patients with a favorable prognosis than the manual approach (Supplementary Figure S4).

These findings indicate that automated TIL assessment using DL on H&E slides could provide prognostic utility for stratifying patients with distinct PFS outcomes.

### Prognostic value of IHC marker expressions

To examine the prognostic value of IHC markers in comparison to H&E-based subgroups, we excluded samples with insufficient registration between H&E and IHC; 72–78 primary tumors and 41–60 pN+ lymph nodes were included for each marker (Supplementary Table S1). Prior to the integration of H&E and IHC data, an examination of the automated protein expression quantification within the tissue and its impact on PFS was conducted through KM analysis. We employed a DL approach to detect, segment, and quantify the number of cells positive for each protein marker, as described in the Methods section [19]. The DL model was applied to the entire IHC image to automatically identify and delineate cells expressing the specific protein marker. Following cell quantification, we categorized the patients into high- or low-expression subgroups based on the median threshold of the number of cells positive for a specific marker of interest within the IHC data.

Independent quantification of IHC expression showed distinct prognostic implications for subgroups defined by high or low protein expression levels. Specifically, high CD4, CD8, and CD20 expression or low ER-beta expression in primary tumors was associated with significantly improved PFS (log-rank test p < 0.05). In addition, high expression of CD3, CD4, CD20, and CD45 in pN+ lymph nodes was also significantly correlated with improved PFS outcomes (log-rank test p-value < 0.05) (Supplementary Figure S5). While high CD4 or CD20 expression was positively associated with PFS in both primary tumors and pN+ lymph nodes, other markers demonstrated tissue-specific associations.

Additionally, the Chi-squared test revealed a significant association between patient status (categorized as ‘progressors’ and ‘non-progressors’) and IHC marker expression of CD4 or CD20 for primary tumors and CD3, CD4, or CD45 for pN+ lymph nodes (Supplementary Figure S3). Again, CD4 levels were consistently significant across both tissue types, whereas other markers displayed varying associations, depending on the tissue examined.

### Understanding the prognostic value of CD8 and CD163 through Integrative analysis of H&E and IHC images

To understand functional cell phenotypes and their cellular composition within the TME, we conducted an integrated analysis utilizing image-based registration of all serial sections of IHC images with H&E staining (Figure 1B). This analysis enabled the incorporation of specific protein markers from the IHC images into the existing H&E-based immune subtypes, thereby further subcategorizing these immune subtypes. We found distinct PFS associations within these immune subtype subcategories across both primary tumors and pN+ lymph node samples.

We hypothesized that we can further stratify immune subtypes identified by the H&E-based DL approach by incorporating 11 proteins from adjacent IHCs. Specifically, we hypothesized that the functional characteristics of TILs and immunosuppressive TME could provide better prognostication. We first characterized the TME by focusing on CD8+ cells and used this information to stratify the IN subgroup into CD8+ high and low patient subgroups in primary tumors and pN+ lymph nodes. As expected, KM analysis showed that the IN CD8+ high patient subgroup consistently showed a better prognosis compared to IN CD8+ low and the rest of the immune subtypes (log-rank test p-value, 0.004) (Figure 3A). Similarly, we used the immunosuppressive M2 macrophage marker, CD163, to define immunosuppressive TME subtypes. We stratified the IN subgroup into CD163+ high and low subgroups for primary tumors and pN+ lymph nodes. KM analysis showed that the IN subgroup with low CD163+ cells (favorable immune microenvironment) showed better PFS in primary tumors compared to the IN subgroup with high CD163+ cells and other subgroups. Conversely, in pN+ lymph nodes, the IN subgroup with high CD163+ cells showed a better PFS outcome than the rest of the subgroups, indicating the tissue-dependent prognostic role of CD163+ cells. A multivariate Cox analysis including disease stage, age, smoking status, ACE-27 score, and sex was conducted to evaluate the prognostic significance of CD8+ and CD163+ immune subtypes within the IN subgroup and other immune subgroups. Our analysis demonstrated that primary tumor subgroups in the IN phenotype with high CD8 (HR, 0.15; 95% CI, 0.043-0.49, p-value = 0.002) and low CD163 (HR, 0.16; 95% CI, 0.045-0.55, p-value = 0.004) had significantly better prognosis compared to other subgroups. In the pN+ lymph nodes, high CD8 (HR, 0.072; 95% CI, 0.010-0.51, p-value = 0.008) and high CD163 (HR, 0.10; 95% CI, 0.020-0.51, p-value = subtypes were associated with improved prognosis (Figure 4).

**Figure 3.**
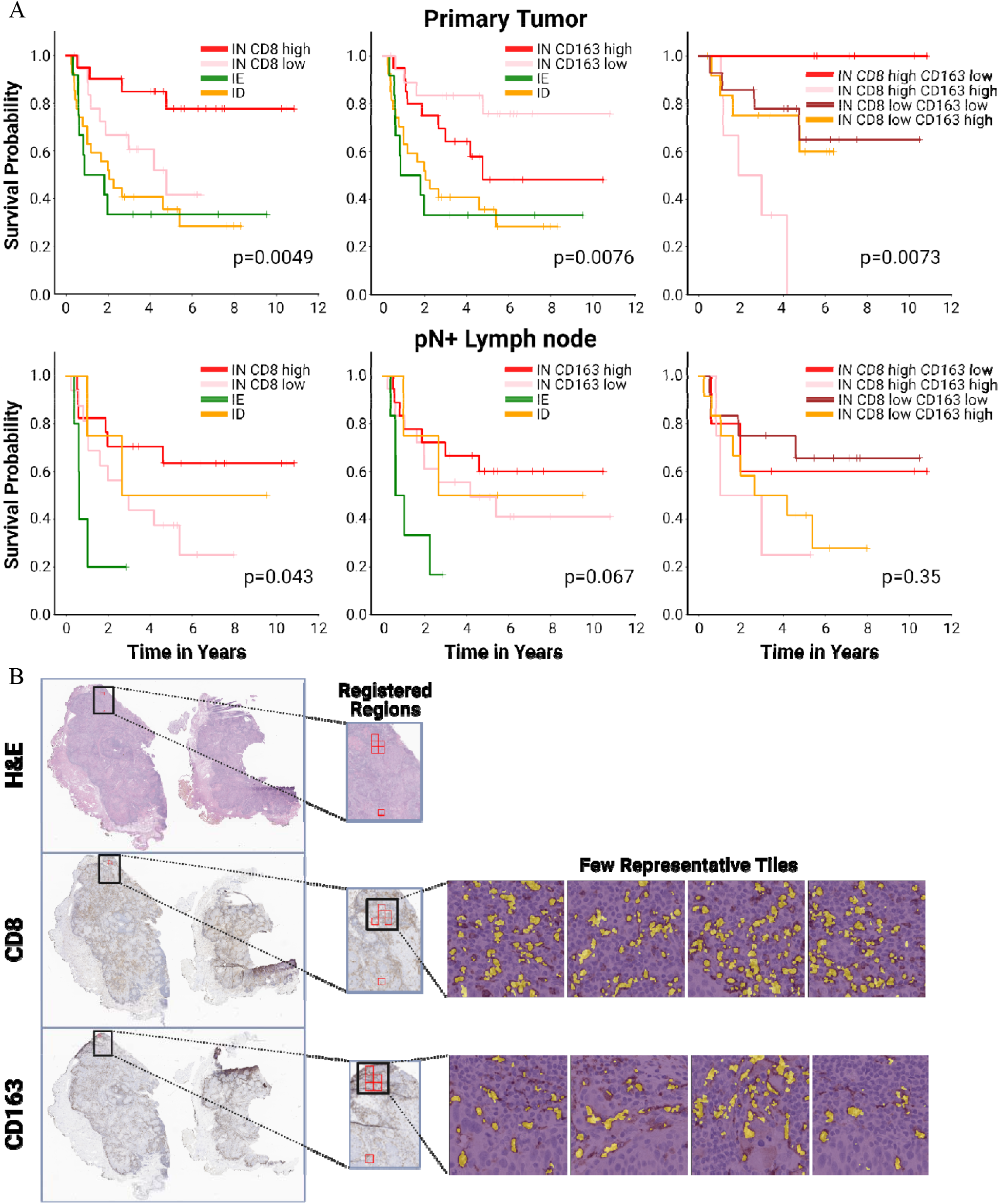
(A) Kaplan-Meier analysis of progression-free survival based on the integration of CD8 and CD163 expression levels along with the identified immune phenotypes from H&E for the primary tumor (top row) and pN+ lymph node (bottom row); IN, immune inflamed; IE, immune excluded; ID immune desert. (B) Visual overview of the organization of the CD8 and CD163 positive cells in the randomly selected registered regions of the tissue for immune inflamed CD8 high and CD163 low patient sample.

**Figure 4.**
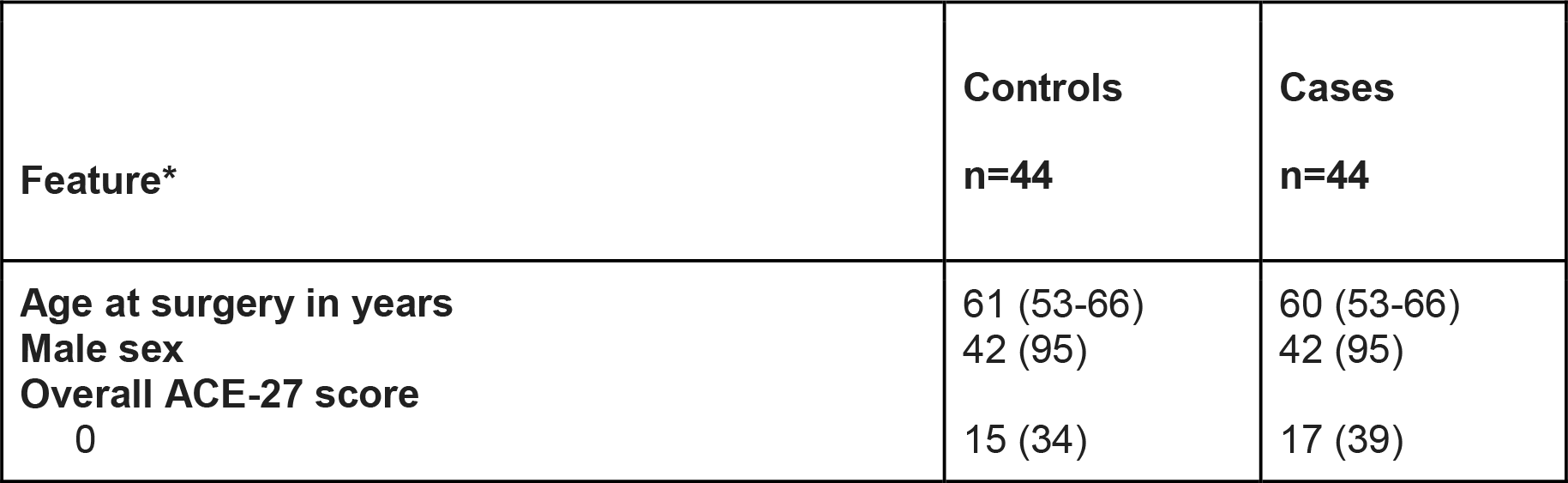
Multivariate Cox analysis, including tumor stage, age, smoking status, treatment management, ACE-27 score, and sex with predicted immune subtypes for primary tumor (top row) and pN+ lymph node (bottom row). IN, immune inflamed; ID, immune desert; IE, immune excluded. S, surgery; R, radiation; C, Chemotherapy.

Finally, to investigate whether a higher level of CD8+ cells, indicative of a more favorable immune microenvironment, coupled with a lower level of CD163+ M2 macrophages, suggestive of a less immunosuppressive TME, correlates with improved PFS, we stratified the IN subgroup using both CD8 and CD163 markers. Interestingly, in the primary tumor, the IN subgroup characterized by high CD8+ and low CD163+ cells showed significantly better prognosis than the IN high CD8+ and high CD163 cell subgroups (univariate KM analysis, log-rank test p = 0.007) (Figure 3A). In the pN+ lymph nodes, the IN subgroup was further characterized by high levels of both CD8+ and CD163+ cells, demonstrating improved prognosis in terms of PFS (univariate KM analysis, log rank test p-value, 0.35) (Figure 3A).

We performed further immune subgroup analyses using the remaining protein markers. In the primary tumor, the IN subgroup, characterized by high expression levels of CD3, CD4, CD20, FoxP3, and ER-alpha and low expression levels of CD45, ER-Beta, PD-L1, and KRT AE1AE3, was associated with favorable PFS (Supplementary Table S3 and S4). In the pN+ lymph nodes, the IN subgroups with high levels of CD8, CD45, and KRT AE1AE3 showed improved PFS (Supplementary Tables S3 and S5). Discrepancies were observed in the enrichment of immune subgroups based on specific protein markers between the primary tumor and pN+ lymph nodes. The distribution of patient subgroups across primary tumors and pN+ lymph nodes and their agreement on different subgroups were compared in a cross-table format (Supplementary Table S2).

Taken together, these findings highlight the importance of considering both H&E-based subtypes and IHC marker expression levels for a comprehensive understanding of the immune landscape within the TME from both primary tumors and pN+ lymph nodes and its implications for patient prognosis.

## Discussion

Head and Neck Squamous Cell Carcinoma is a heterogeneous cancer arising from the oral cavity, pharynx, larynx, and other related anatomical sites [20], and is usually linked to tobacco and alcohol consumption [21]. However, in recent years, a distinct subgroup of HNSCC associated with HPV infection has emerged as a distinct clinical factor. HPV(+) OPSCC is characterized by unique clinical and molecular features, and the TME plays an important role in the clinical course and treatment response of these tumors.

Previous studies have shown that HPV(+) OPSCC is associated with a more favorable prognosis than its HPV(-) counterpart [22, 23]. This improved outcome was partly attributed to the enhanced immune response within the TME. HPV(+) OPSCC tumors often exhibit increased infiltration of cytotoxic T lymphocytes (CTLs) and higher levels of immune checkpoint proteins, such as PD-L1 [24]. This immune-active TME has paved the way for the successful use of immune checkpoint inhibitors (ICIs) in the treatment of HPV(+) HNSCC [25,26,27]. Unfortunately, only 15–20% of patients with HNSCC benefit from ICI, and this poor outcome is increasingly ascribed to the peculiar characteristics of the TIME [28]. Understanding TME is crucial for predicting the response to immunotherapy, optimizing treatment regimens, and identifying biomarkers that can guide patient selection.

In this study, we investigated the prognostic value of the colocalization of TILs with tumor and stroma regions using H&E staining combined with functional characterization of tumor and immune cells from IHC stains in a matched case-control study of patients with HPV(+) OPSCC. We leveraged DL approaches for an integrative analysis of both H&E- and IHC-stained slides and demonstrated the categorization of H&E slides into distinct immune subtypes using an AI algorithm. This integrative approach not only allowed us to overcome some of the limitations associated with H&E staining alone but also enabled us to reveal the functional phenotypes of cells within the TME. In particular, we focused on the cytotoxic and less immune-suppressive TME, which is characterized by the presence of CD8+ and CD163+ cells. This in-depth TME analysis led to the identification of specific immune subgroups that are significantly associated with PFS. Moreover, our findings provided the immunological landscape and prognostic relevance of the TME in the primary tumor and pN+ lymph node.

While the primary focus of this work was on TILs co-organized with CD8+ and CD163+ cells within the TME, we also extended our analysis to other protein markers related to epithelial cells, B cells, and regulatory T cells. For example, we found that the IN subgroups enriched in FoxP3+ cells in primary tumors were associated with improved PFS (log-rank test, p = 0.008) (Supplementary Figure S6 (A, B)). Although FoxP3+ cells are known for their immunosuppressive function, our findings suggest a unique role for FoxP3 in HPV(+) OPSCC. Recent studies have shown that similar findings related to FoxP3+ cells correlate with improved prognosis in HPV(+) OPSCC [29,30]. Other interesting observations include the association of high levels of CD20+ B cells and low levels of KRT AE1AE3+ epithelial cells with improved PFS in primary tumors, but not in pN+ lymph nodes (Supplementary Figure S6 (A, B)). These results confirm the tissue-dependent prognostic role of various cell types, including CD20+ B cells, KRT AE1AE3+ epithelial cells, and FoxP3 Tregs, in the TME in primary tumors and pN+ lymph nodes. Although the implications of the presence of other cell phenotypes associated with PFS are intriguing, they point toward potential opportunities for future research.

This study has several limitations. For instance, while we employed serial adjacent sections of H&E and IHCs from a single tumor and matched pN+ lymph nodes, some slides were situated distally, and the tissue alignment was not always perfect. In addition, potential discrepancies in TME characterization could be introduced when comparing distal slides. These challenges can be addressed using multiplex IHC, spatial proteomics, or In Situ Molecular Imaging technologies, which enable the staining of multiple proteins on the same slide. Another limitation is the small patient cohort, which restricted our analysis to a limited set of protein markers. Future studies with larger datasets could resolve this issue. Finally, the underlying molecular mechanisms driving the prognostic relevance of these specific TME subgroups remain to be explored further.

In summary, our study highlights the utility of integrating histological and immunohistochemical analyses, paired with deep learning techniques, for the comprehensive characterization of TME functional phenotypes. This approach offers the capability to leverage information from TME to stratify patients more effectively into subgroups with distinct survival outcomes, thereby advancing the field of precision oncology care for patients with HPV+ OPSCC.

## ACKNOWLEDGEMENTS

This study was funded internally at Mayo Clinic, Florida. We thank the patients and their families.

## Notes

**CONFLICT OF INTEREST:** None

### Competing Interest Statement

The authors have declared no competing interest.

