## Supplementary figures and images for "Integrative Spatial Analysis of H&E and IHC Images Identifies Prognostic Immune Subtypes Correlated with Progression-Free Survival in Human Papillomavirus (HPV)-Related Oropharyngeal Squamous Cell Carcinoma"

### Figure S1

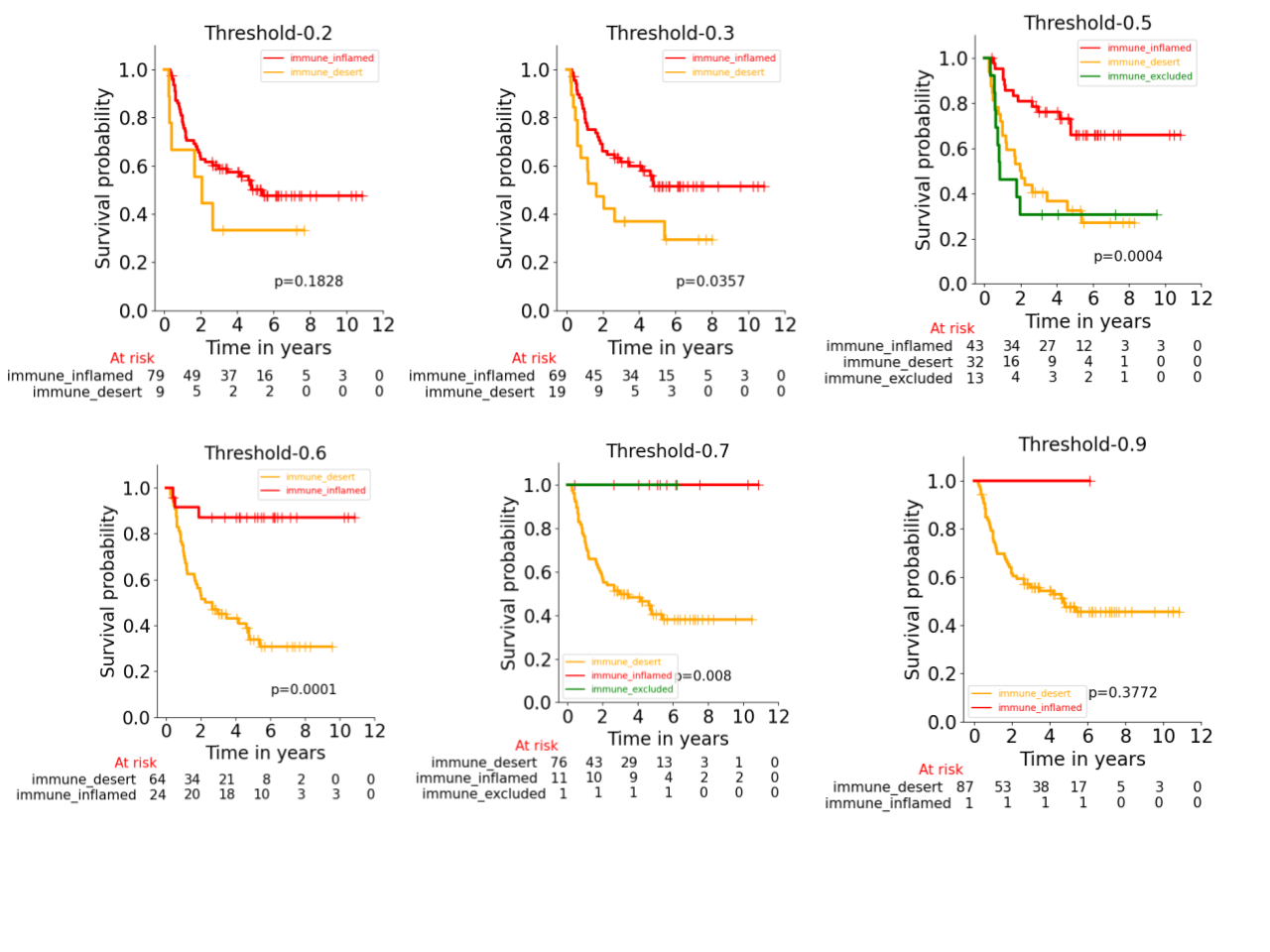

### Figure S2

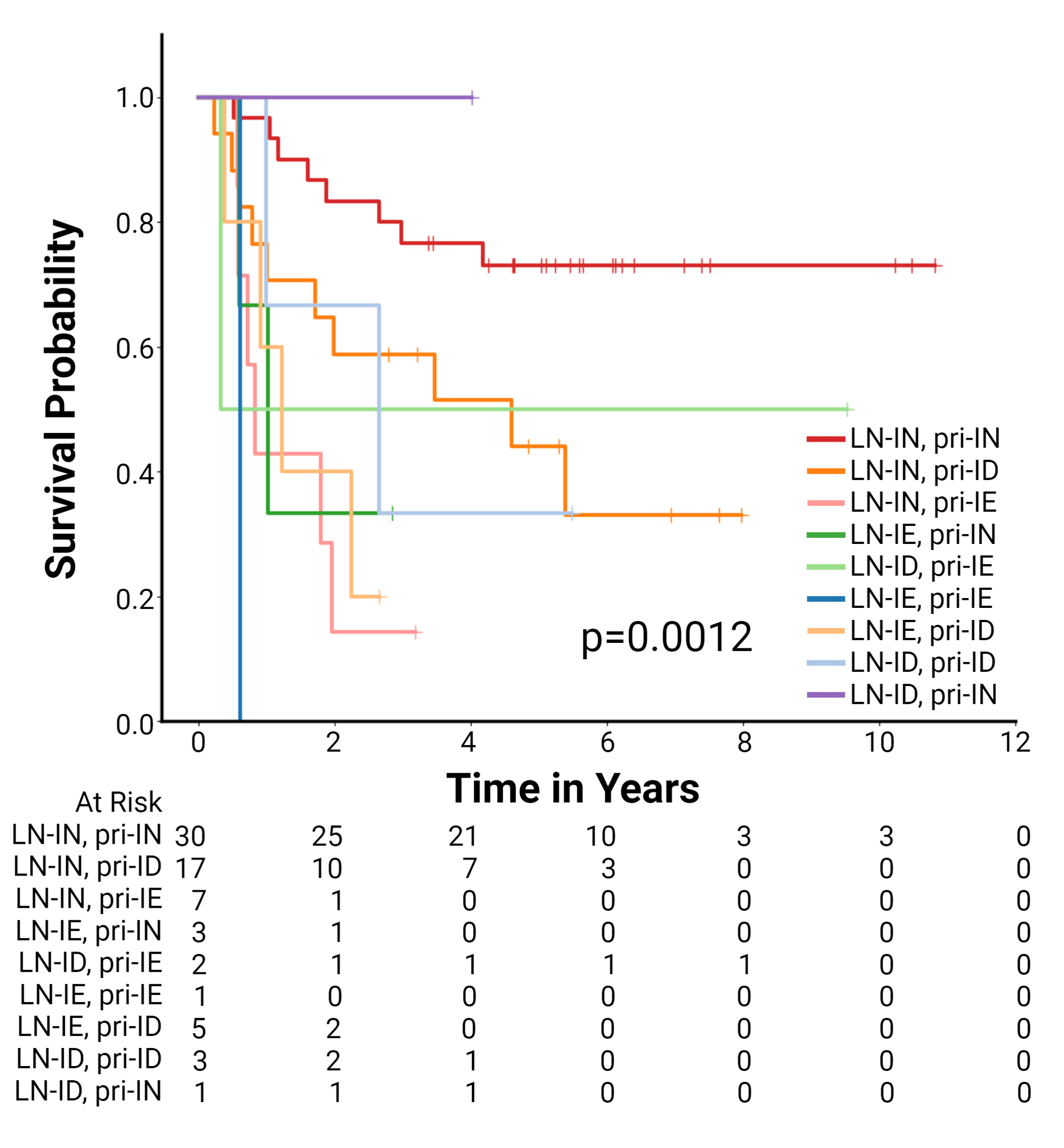

### Figure S3 A

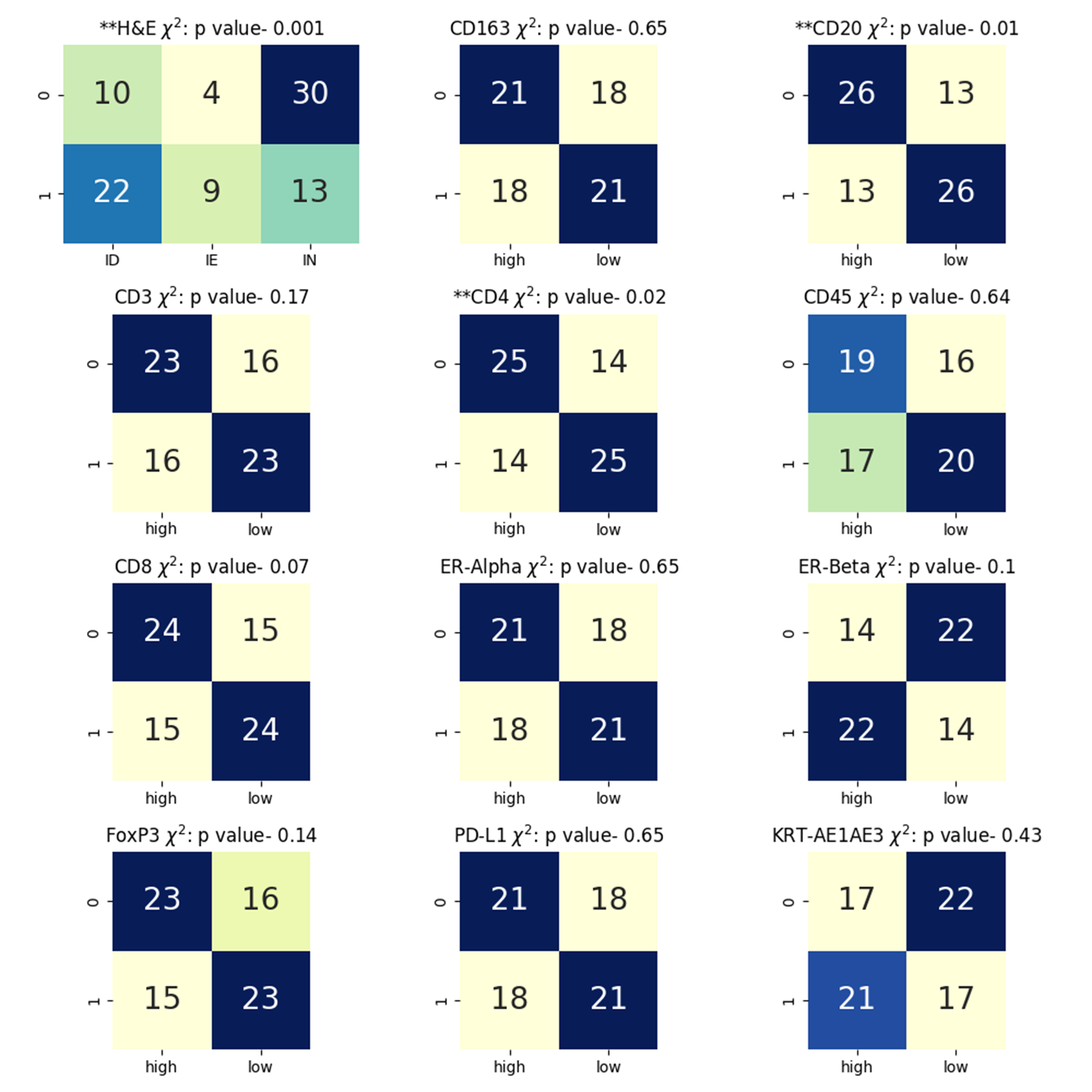

### Figure S3 B

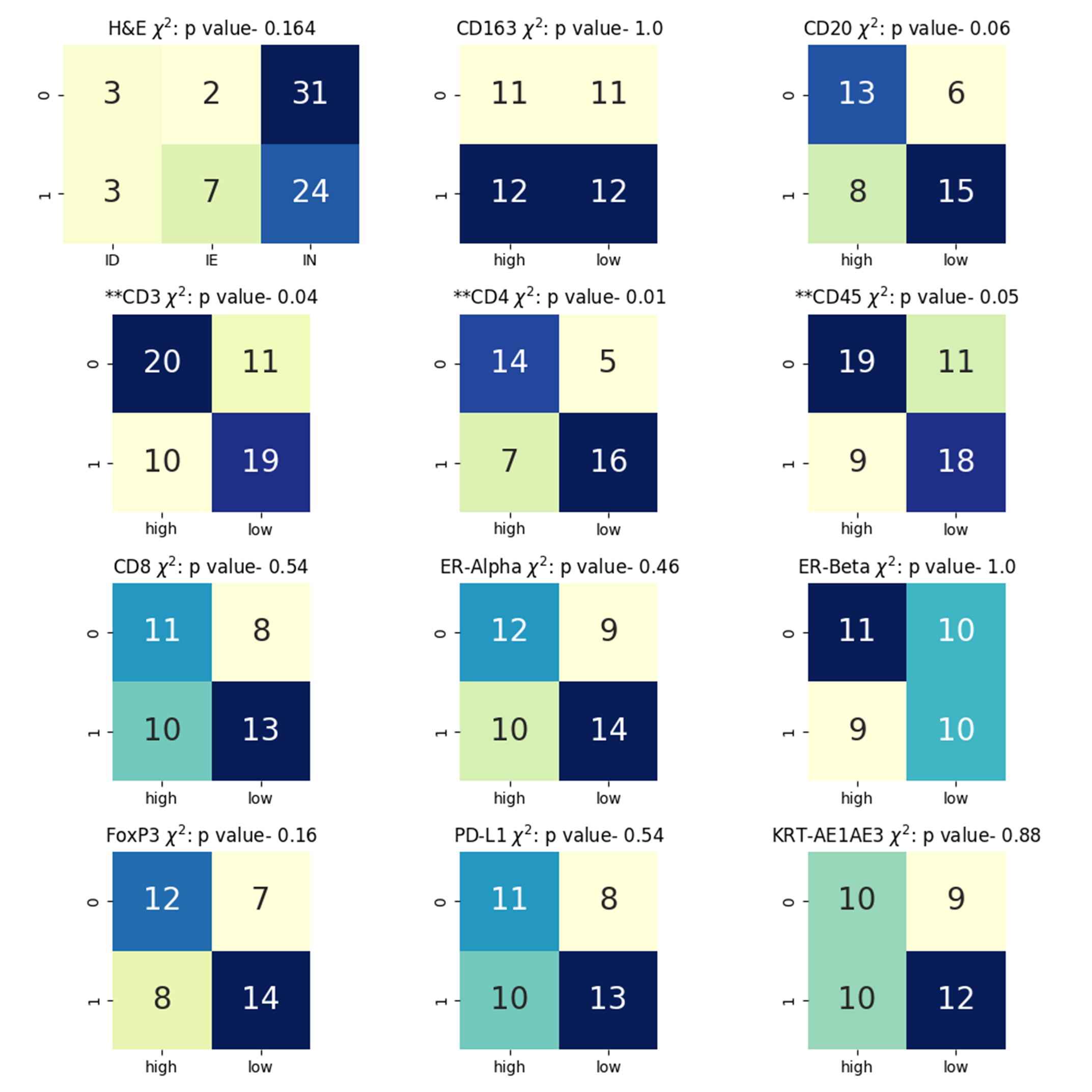

### Figure S4

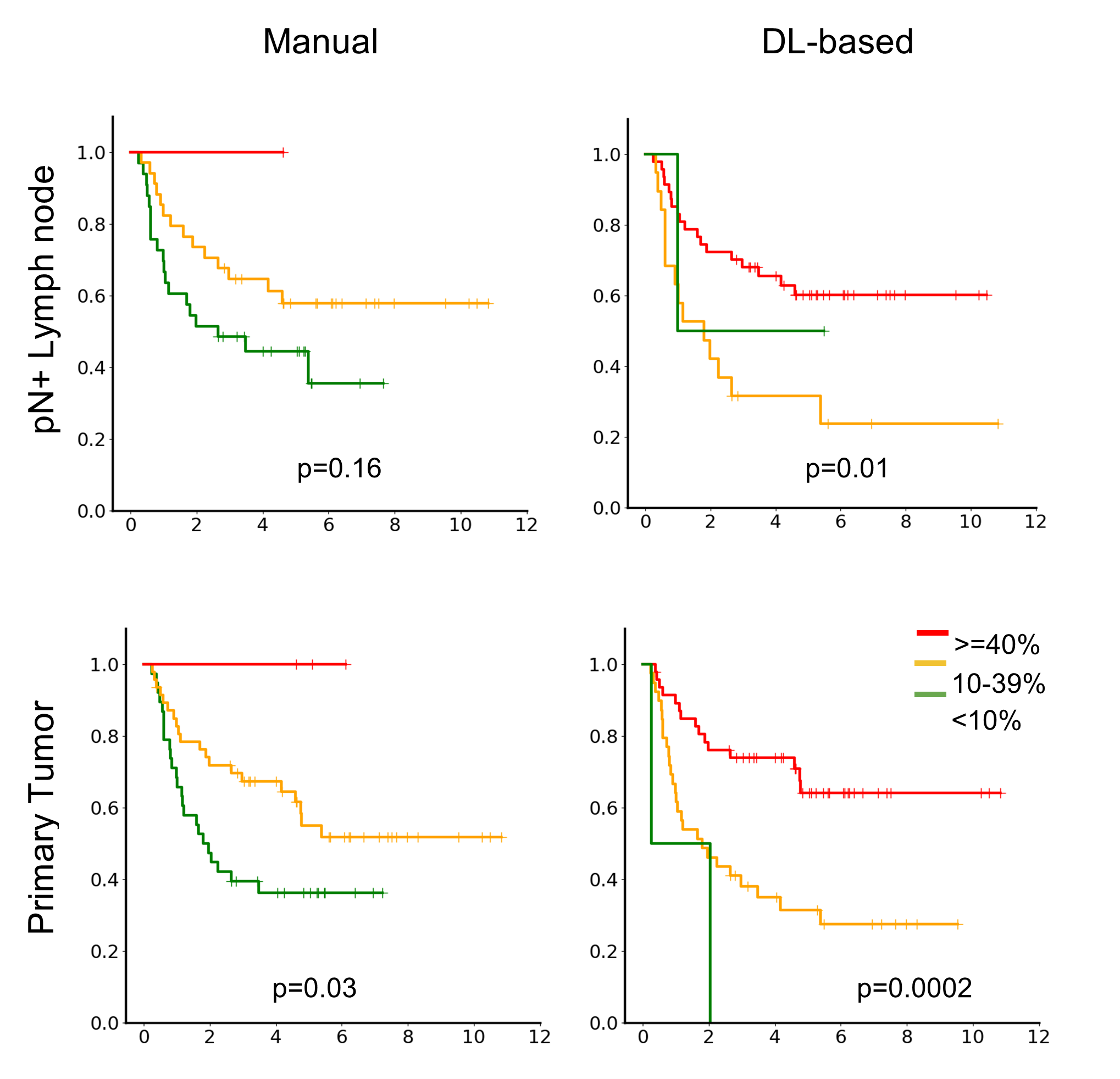

### Figure S5 A

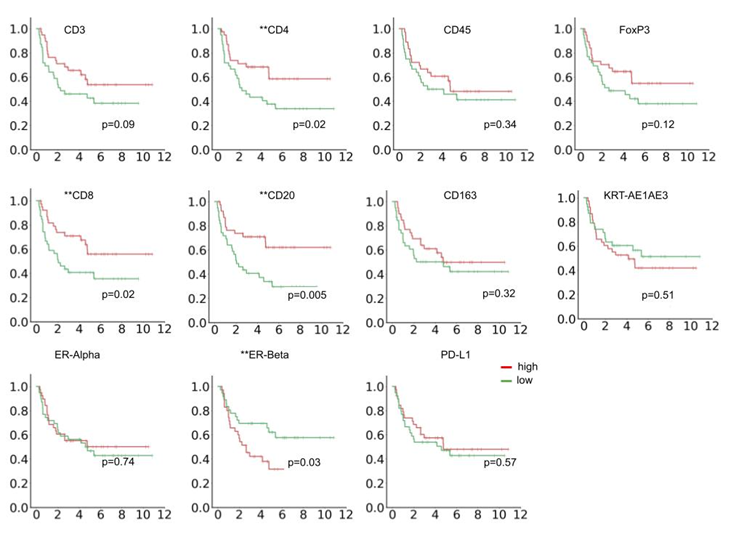

### Figure S5 B

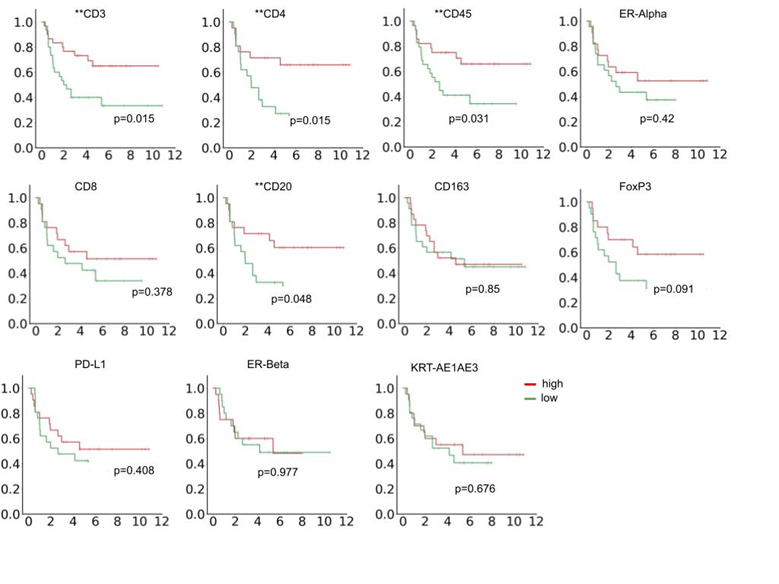

### Figure S6 A

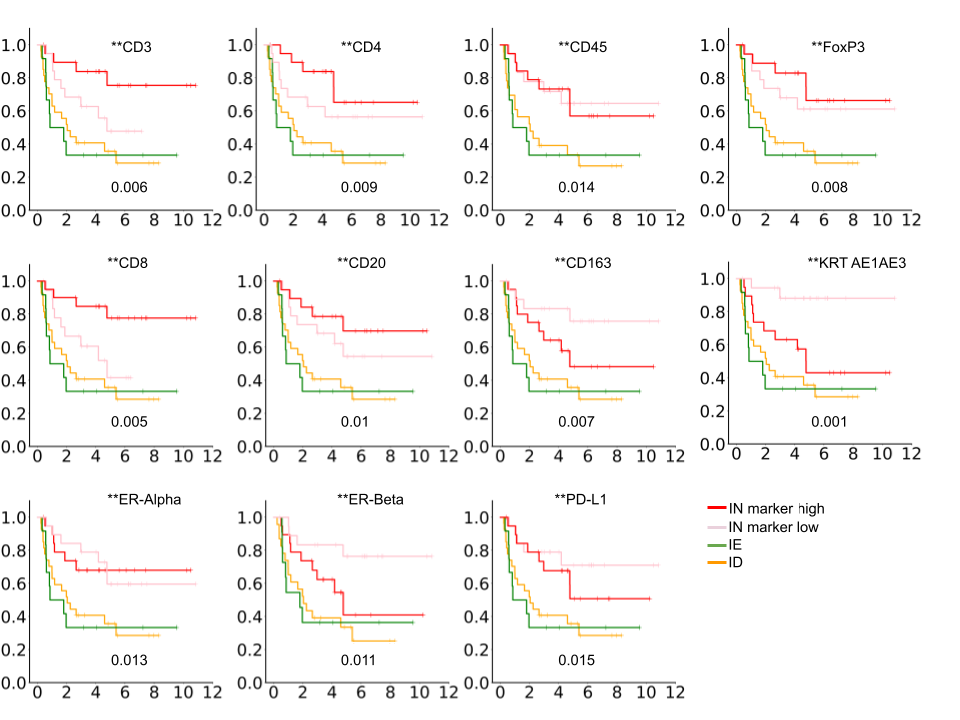

### Figure S6 B

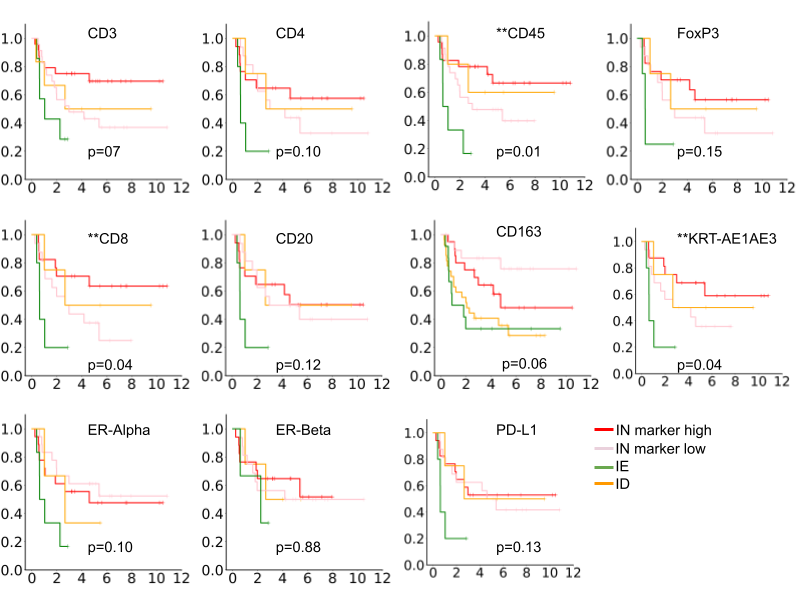
